# A single amino acid substitution in MdLAZY1A dominantly impairs shoot gravitropism in *Malus*

**DOI:** 10.1101/2023.04.05.535771

**Authors:** Laura Dougherty, Ewa Borejsza-Wysocka, Alexandre Miaule, Ping Wang, Desen Zheng, Michael Jansen, Susan Brown, Miguel Piñeros, Christopher Dardick, Kenong Xu

## Abstract

Plant architecture is one of the most important factors that determines crop yield potential and productivity. In apple (*Malus*), genetic improvement of tree architecture has been challenging due to a long juvenile phase and their growth as complex trees composed of a distinct scion and a rootstock. To better understand the genetic control of apple tree architecture, the dominant weeping growth phenotype was investigated. We report the identification of *MdLAZY1A* (MD13G1122400) as the genetic determinant underpinning the *Weeping* (*W)* locus that largely controls weeping growth in *Malus*. *MdLAZY1A* is one of the four paralogs in apple that are most closely related to *AtLAZY1* involved in gravitropism in *Arabidopsis*. The weeping allele (*MdLAZY1A-W*) contains a single nucleotide mutation c.584T>C that leads to a leucine to proline (L195P) substitution within a predicted transmembrane domain that co-localizes with Region III, one of the five conserved regions in LAZY1-like proteins. Subcellular localization revealed that MdLAZY1A localizes to the plasma membrane and nucleus in plant cells. Over-expressing the weeping allele in apple cultivar Royal Gala (RG) with standard growth habit impaired its gravitropic response and altered the growth to weeping-like. Suppressing the standard allele (*MdLAZY1A-S*) by RNA interference (RNAi) in RG similarly changed the branch growth direction to downward. Overall, the L195P mutation in MdLAZY1A is genetically causal for weeping growth, underscoring not only the crucial roles of residue L195 and Region III in MdLAZY1A-mediated gravitropic response, but also a potential DNA base editing target for tree architecture improvement in *Malus* and other crops.

## Introduction

Plant form or architecture is one of the most important factors that determines crop yield potential and productivity. Genetic improvement of plant architecture in cereal crops led to a dramatic increase in grain yield, a revolutionary accomplishment in agriculture known as ‘the Green Revolution’ (Evenson and Gollin, 2003). As such, a tremendous amount of research has been devoted to understanding the underlying biological processes that dictate plant architecture (Teichmann and Muhr, 2015; Wang et al., 2018). Recent advances in understanding how gravity affects plant form are of great importance in plant biology and genetic improvement as gravity is a constant natural physical force imposed on all living organisms (Takahashi et al., 2021).

The term ‘gravitropism’ refers to growth response of a cell or organism to gravity. In plants, shoots exhibit negative gravitropism growing upward against gravity, while roots show positive gravitropism growing downward in the direction of gravity. However, most plant lateral organs, such as branches and leaves grow at a certain angle in relation to gravity termed gravitropic set-point angle (GSA) (Digby and Firn, 1995). Although different plant organs, such as roots, hypocotyls, and inflorescences may differ in the molecular mechanism of their gravitropic responses (Chen et al., 1999; Tasaka et al., 1999; Baldwin et al., 2013), the gravitropic pathway generally comprises three sequential steps: perception, biochemical signaling and differential growth (Nakamura et al., 2019; Vandenbrink and Kiss, 2019; Takahashi et al., 2021). Perception of gravity occurs in shoot endodermal cells and root columella cells that are collectively referred to gravity sensing cells or statocytes (Sack, 1991; Fukaki et al., 1998; Morita, 2010; Vandenbrink and Kiss, 2019). An important characteristic of statocytes is that they contain many statoliths starch granule-filled amyloplasts. When plants are reoriented, the amyloplasts sediment to the new lower side of statocytes, thereby converting the potential energy of gravity into an initial intracellular signal in the elongation zone (Kiss et al., 1989). Once the signal generated by amyloplast sedimentation is perceived, an unknown mechanism results in a rapid polarization of the LAZY1-like proteins to the plasma membrane on the lower side of statocytes, leading to a cascade of polar recruitment of RCC1-like domain (RLD) proteins from cytoplasm and polar re-localization of auxin efflux carrier proteins (Furutani et al., 2020), such as such as PIN3 (Friml et al., 2002; Kleine-Vehn et al., 2010). Consequently, the direction of auxin flow is reoriented downward, and a polarized gradient of auxin is created, leading to differential elongation growth on opposing sides of the root or shoot (Vandenbrink and Kiss, 2019; Takahashi et al., 2021).

The *LAZY1* gene was first identified in rice as a plant-specific gene involved in gravitropic response, and its loss-of-function or null alleles are genetically responsible for the prostrate growth phenotype of ‘lazy’ rice lines that have partially impaired gravitropism (Li et al., 2007; Yoshihara and Iino, 2007). Subsequently, orthologs of *LAZY1* were found to play a similar role in regulating gravitropic response and/or branch angle in *Arabidopsis* (Yoshihara et al., 2013), *Zea mays* (Dong et al., 2013), and *Populus* (Xu et al., 2017). The AtLAZY1 protein has five conserved regions (Yoshihara et al., 2013) with Region II containing a highly conserved ‘IGT’ motif that typifies a family of proteins to include *TILLER ANGLE CONTROL 1 (TAC1)* (Dardick et al., 2013; Waite and Dardick, 2021). The indispensable role of amino acids ‘IGT’ in LAZY1 function was demonstrated clearly in a recessive rice mutation G74V (Chen et al., 2022). Site-directed mutagenesis-based amino acid substitutions in all five conserved regions of AtLAZY1 revealed that Region I is required for plasma membrane localization, Regions II and V are critical to its function; and Regions III and IV are tolerant to the amino acid substitutions studied (Yoshihara and Spalding, 2020).

In apple (*Malus*), genetic improvement of tree architecture has been challenging due to a long juvenile phase and their growth as complex trees composed of a distinct scion and a rootstock. Another limiting factor is that the genetic control of apple tree forms remains poorly understood. Currently, apple tree forms are grouped into four ideotypes (I-IV): columnar (as represented by Wijcik McIntosh), spur (Starkrimson), standard (Golden Delicious), and weeping (Granny Smith and Rome Beauty) based on the tree overall growth and fruiting patterns (Lespinasse and Delort, 1986; Lespinasse, 1992; Costes et al., 2006; Pereira-Lorenzo et al., 2009; Höfer et al., 2013). However, the drooping branches in ideotype IV cultivars are caused by their tip-bearing fruit, which is distinctive from the weeping phenotype caused by the natural downward growth of branches. Weeping cultivars are rare in cultivated apples (Lindén and Iwarsson, 2014), but common in crabapples for ornamental and landscape gardens, such as the popular weeper Red Jade (Brown et al., 2004). These weeping cultivars offer a mutant tree form that can eliminate the need of labor-intensive branch bending (90°-120° from the vertical main trunk) in orchard management (Ferree and Schupp, 2003; Musacchi and Greene, 2017). Additionally, they also present an opportunity to study and understand the underlying mechanisms of apple tree architecture. However, only a few genetic studies were reported, and the understanding is limited to that the weeping phenotype is controlled by a dominant allele *Weeping* (*W)* (Sampson and Cameron, 1965; Brown, 1992; Alston et al., 2000; Just, 2001).

To fill the knowledge gap for improvement of tree architecture, we investigated the weeping phenotype and reported four genomic regions relevant in genetic control of weeping growth, including *W* mapped on chromosome 13 and three other regions on chromosomes 10 (*W2*), 16 (*W3*), and 5 (*W4*), respectively (Dougherty et al., 2018). Among them, the *W* locus had the largest genetic effect on weeping growth, and was localized to a 982-kb genomic region containing 72 predicted genes (Dougherty et al., 2018) in the apple reference genome (Daccord et al., 2017). As part of an ongoing effort, we report that gene MD13G1122400, designated *MdLAZY1A*, one of the four apple orthologs most closely related to *AtLAZY1*, is the genetic determinant underlying *W*. We show that a single amino acid substitution (L195P) in the conserved Region III in the weeping allele is sufficient to disrupt the function of MdLAZY1A in both alleles in heterozygous plants and alter branch growth from standard to weeping-like, underscoring the crucial role of residue L195 and Region III in MdLAZY1A mediated gravitropism in apple.

## Results

### Genetic characterization of the *Weeping* (*W*) locus

To minimize the *W* region (982-kb) defined by SSR markers Ch13-8181, Ch13-8547 and Ch13-9530 on chromosome 13 (Dougherty et al., 2018), the mapping populations were increased from 217 to 1246 progenies in three F_1_ and two open-pollinated populations generated from the weeping cultivars Cheal’s Weeping (CW), Red Jade (RJ) and Louisa (**Supplemental Table S1**). The segregation between the weeping and standard phenotypes fit a 1:1 ratio in the F_1_ populations (p=0.2207-0.6042) as well as in the open-pollinated progenies of CW and RJ (p=0.3685-0.7888) (**Supplemental Table S1**). Genotyping of the 1246 individuals with the *W*-linked marker Ch13-8547 (in *MdLAZY1A’s* promoter region) and/or H_9031 (detects the L195P mutation in MdLAZY1A, **Supplemental Table S2**) demonstrated that the phenotypic segregation was largely determined by genotypes *Ww* (weeping, 513/555 or 92.4%) and *ww* (standard, 544/554 or 98.2%) (**Fig. 1A-D**). However, 42 (7.6%) seedlings of a weeping genotype *Ww* displayed a standard phenotype (**Fig. 1A**). A close look indicated that 37 of the 42 seedlings were from the progeny of CW (**Fig. 1B, Supplemental Table S1**), presumably caused by the three loci *W2-W4* identified in CW (Dougherty et al., 2018).

**Figure 1.**
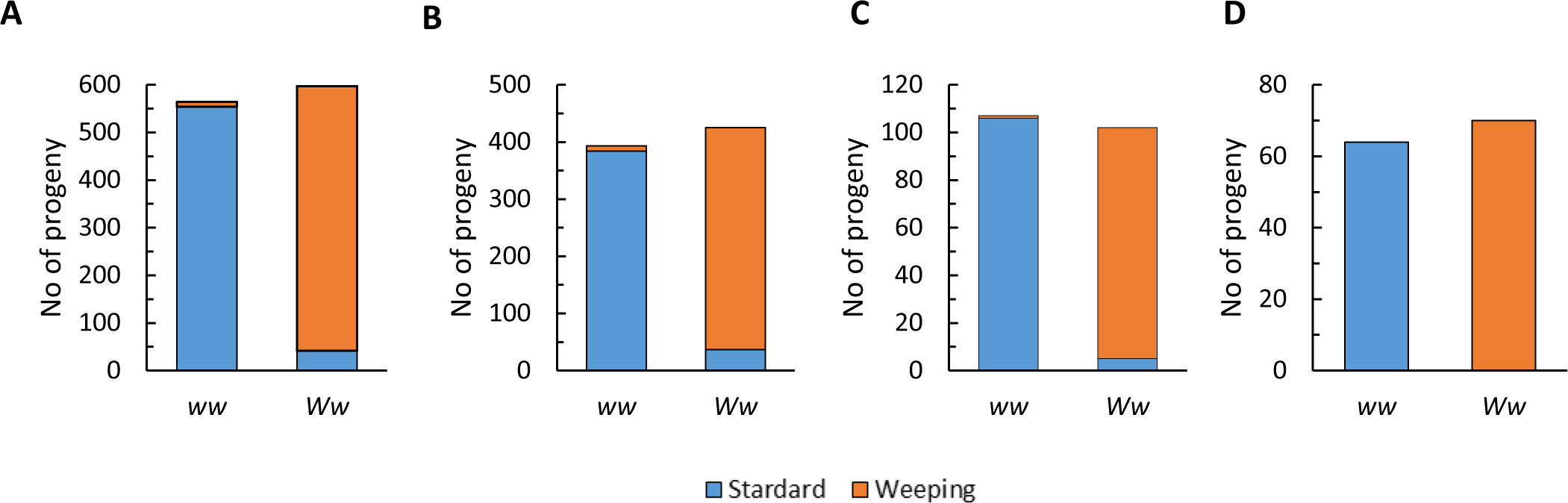
Phenotypic segregation in mapping populations. **A**, All progenies combined (n=1161, p=0.3629). **B-D**, Progenies of weeping cultivars Cheal’s Weeping (B) (n=818, p=0.3662), Red Jade (C) (n=209, p=0.3685), and Louisa (D) (n=134, p=0.6042). There were 85, 57, 22 and 6 seedlings trees excluded in A, B, C, and D due to intermediate phenotype or deaths, respectively. Standard= standard and standard-like phenotypes combined. Weeping= weeping and weeping-like phenotypes combined. Genotypes *WW* and *ww* were determined by *W*-linked markers.

Combined with the single nucleotide variants (SNVs) identified in an RNA-seq analysis (see below), seven high resolution melting (HRM) markers were developed to target SNVs in coding sequences of annotated genes within the *W* region (**Supplemental Table S2**). Based on the genotypes of these markers, 47 informative recombinants were identified, including 23 derived from a recombined gamete between markers H_8694 and H_9574 in the progeny of CW (21) and Louisa (2), and 24 from a recombined gamete between markers H_8735 and Ch13-8547 in the progeny of RJ (**Supplemental Table S1**). A schematic representation of the recombinants by marker genotypes and growth habit showed that *W* co-segregated with markers Ch13-8547 and H_9031 within a 311-kb genomic region between markers H_8898 (at 8,898^th^ kb) and H_9209 (at 9,209^th^ kb) on chromosome 13 (**Supplemental Fig. S1**). The marker genotypes also indicated that cultivars CW, RJ and Louisa are independent sources of weeping (**Supplemental Fig. S1**).

### *MdLAZY1A* Is a strong candidate gene underpinning *W*

There are 23 protein encoding genes annotated in the 311-kb *W*-region in the apple reference genome (Daccord et al., 2017) (**Supplemental Table S3**). To help identify the candidate gene(s) under *W*, an RNA-seq experiment was conducted using actively growing shoot apex tissues sampled from seven weeping and five standard individuals from the families of CW and RJ (**Supplemental Tables S3-S4**). The raw and clean reads generated were 471.3 and 408.2 million in total, and 39.3±15.4 and 34.0±15.8 million per sample on average, respectively. Of the clean reads, 315.2 million (77.6%) were mapped to the apple reference genome, equivalent to 26.3±11.8 million mapped reads per sample (**Supplemental Table S4**). This detected 32,369 genes expressed (reads per kilo base per million mapped reads-RPKM >1.0 in one or more of the 12 RNA-seq samples) non-redundantly, 243 of which were differentially expressed genes (DEGs) (RPKM fold change>1.5, and FDR P<0.05) between weeping and standard in the two families. Under *W*, 20 of the 23 annotated genes were expressed; however, no DEGs were found between weeping and standard, suggesting the candidate genes of *W* would be among the 20 expressed genes (**Supplemental Table S3**).

To further shorten the list of *W* candidate genes, the 12 individual RNA-seq reads-mappings were merged into four groups according to their weeping and standard phenotypes in families CW and RJ, respectively (**Fig. 2A**). DNA variant calling in the four merged RNA-seq sample groups identified 45,789 and 17,169 SNVs specific to weeping in the coding sequences in CW and RJ families, 13,771 and 4,917 of which were non-synonymous, respectively (**Fig. 2A**). Among the weeping specific non-synonymous SNVs, 13,498 were unique to CW, 4,644 unique to RJ, and 273 common to both CW and RJ (**Fig. 2A**). In the *W* region of 311-kb, eight of such weeping specific non-synonymous SNVs were unique to CW, zero to RJ, and two were common to both RJ and CW (**Fig. 2A-E, Supplemental Table S5**).

**Figure 2.**
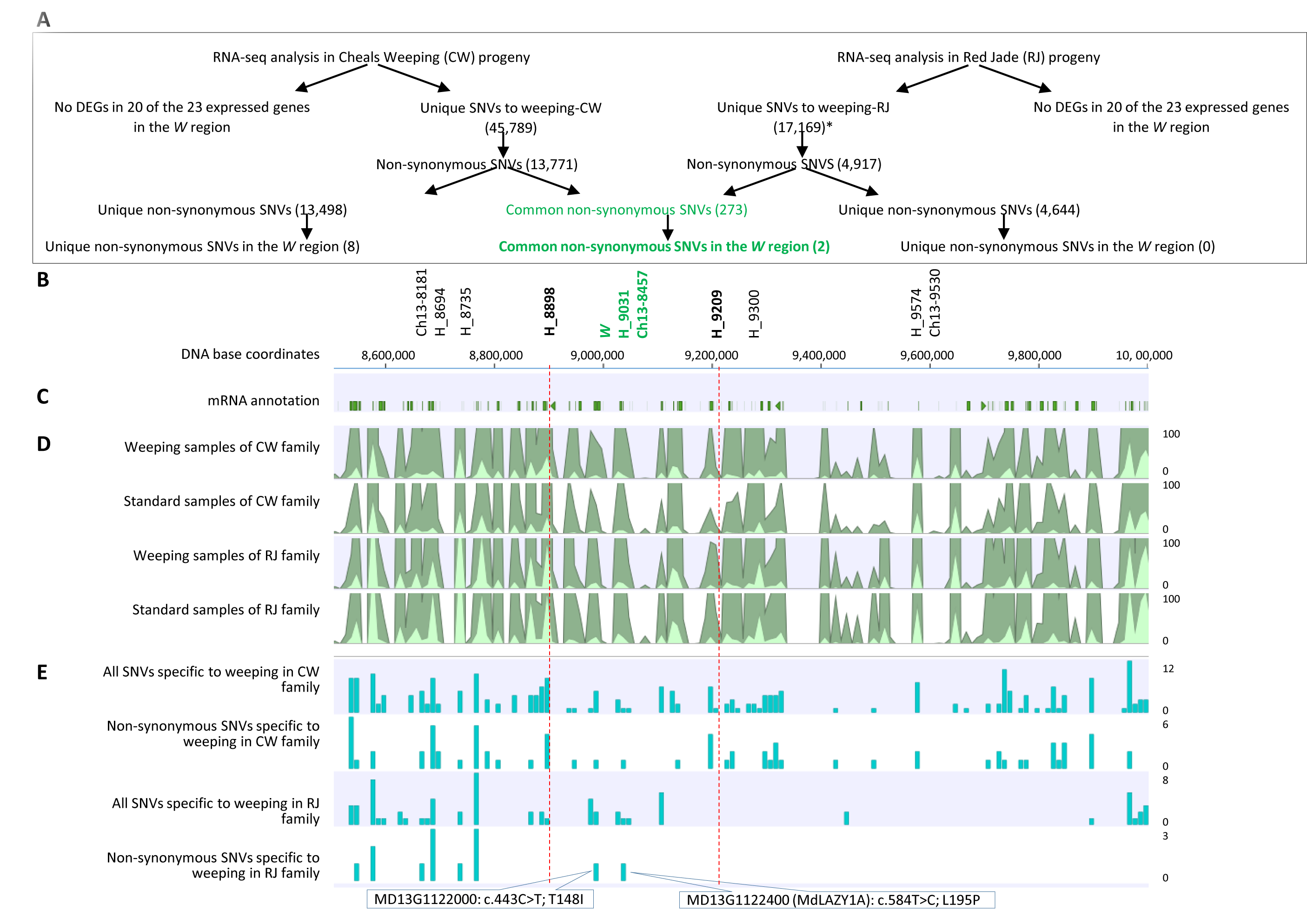
RNA-seq based identification of non-synonymous SNVs specific to weeping phenotype in Cheal’s Weeping (CW) and Red Jade (RJ) families. **A**, A flowchart illustrating the steps taken to identify non-synonymous SNVs specific to weeping in the two families. **B**, DNA markers and their physical locations in the *W* region of 1.5 (8.5-10.0) Mb on chromosome 13 in the apple reference genome. **C**, Diagram of mRNA models annotated. **D**, A compact view of RNA-seq reads coverage (shown 0-100x) in the merged weeping and standard samples from the CW and RJ families. **E**, A compact view of all and non-synonymous SNVs identified specific to the merged weeping samples in the CW and RJ families. The two non-synonymous SNVs that are specific to weeping phenotype in both CW and RJ families are indicated. The 311-kb *W* region between markers H_8898 and H_9209 is indicated by dotted lines. The numbers on the right indicate the scale of coverage.

Between the two weeping specific non-synonymous SNVs common to both CW and RJ, one was the nucleotide mutation (Chr13_8,985,133) from AC_443_T to AT_443_T (c.433C>T) in the open reading frame (ORF) of gene MD13G1122000, leading to a threonine to isoleucine substitution T148I (**Fig. 2E, Supplemental Fig. S2A; Supplemental Table S5**). MD13G1122000 contains a domain of unknown function 1664 (DUF1664) and is most closely related to AT1G25510 (eukaryotic aspartyl protease family protein) in *Arabidopsis*. The other was a base mutation (Chr13_9,031,158) from CT_584_T to CC_584_T (c.584T>C) in the ORF of MD13G1122400, resulting in a leucine to proline substitution L195P (**Fig. 2E; Supplemental Fig. S2B; Supplemental Table S5**). MD13G1122400, designated *MdLAZY1A* encoding a protein of 384 amino acids, is one of the four counterparts of *Arabidopsis AtLAZY1* (AT5G14090) in apple. Given the role of *LAZY1*-like genes in regulation of gravitropism in plants (Li et al., 2007; Yoshihara and Iino, 2007; Yoshihara et al., 2013), *MdLAZY1A* was considered a strong candidate gene underpinning *W* controlling weeping growth.

### Mutation L195P in MdLAZY1A is a signature of weeping cultivars in *Malus*

To understand the difference between *MdLAZY1A’s* weeping (*MdLAZY1A-W*) and standard (*MdLAZY1A-S*) alleles, their genomic sequences were determined in three weeping cultivars RJ, CW, and Louisa and two standard cultivar/selections Golden Delicious and NY-051, leading to seven standard (containing CT_584_T) and three weeping (carrying CC_584_T) alleles. Sequence alignment uncovered important changes in both the promoter and the coding sequences in addition to nucleotide changes in non-coding and non-transcribed sequences (**Fig. 3A-C, Supplemental Fig. S3**). In the coding sequences, which were also confirmed by sequencing cDNA clones from CW (**Supplemental Fig. S4**), 29 SNVs were revealed across the ten alleles (**Supplemental Fig. S3**), 15 of which were non-synonymous (**Supplemental Fig. S4**). However, 11 of the 15 non-synonymous SNVs were observed as a single event, including eight in NY-051-S2 (standard allele 2), two in Louisa-S1 and one in Louisa-W (weeping allele), suggesting these 11 SNVs were unlikely to play a role in weeping. The remaining four non-synonymous SNVs were present in three or more alleles; however, mutation c.584T>C was the only one that was specific to the three weeping cultivars, while the other three SNVs were present in both weeping and standard alleles (**Supplemental Fig. S4**). Direct sequencing of eight additional weeping cultivars (**Fig. 3D**), together with genome sequence curation in 28 other standard *Malus* accessions (**Supplemental Table S6)** (Duan et al., 2017) available in NCBI-SRA databases confirmed that the mutation c.584T>C was associated with weeping exclusively, further supporting that the L195P (c.584T>C) mutation in MdLAZY1A was specific to weeping.

**Figure 3.**
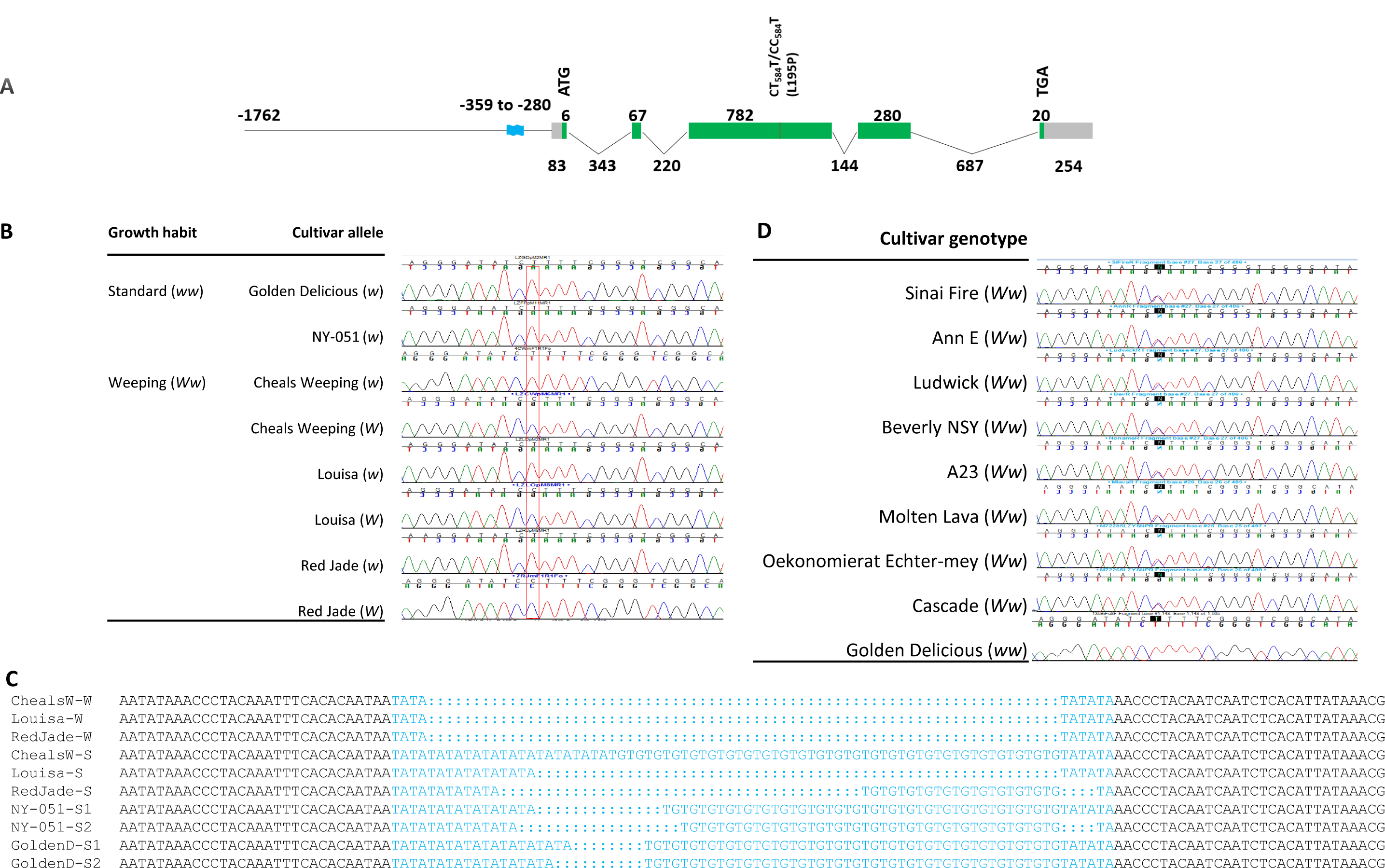
Genomic sequence characterization of *MdLAZY1A* alleles. **A,** Diagram of *MdLAZY1A* genomic structure. Straight line: promoter region. Wave thick line in blue: SSR region. Vertical short black line: mutation c.584T>C, i.e. CT_584_T to CC_584_T leading to L195P. Reflected lines: introns. Green filled bars: coding sequences. Grey filled bars: untranslated regions. ATG: start codon. TGA: stop codon. The numbers indicate sizes (bp) of corresponding segments. The negative numbers refer to positions in the promoter region in relation to the ATG site. **B,** Sequence chromatogram of genomic DNA clones from standard apples Golden Delicious and NY-051 and weeping cultivars Cheal’s Weeping, Louisa and Red Jade. The weeping and standard distinctive SNV at base 584 is indicated. **C,** Sequence alignment of the promoter SSR region in MdLAZY1A weeping (W) and standard (S) alleles. Bases in blue stand for the complex SSR region ‘(TA)_2-12_(TG)_0-25_(TA)_1-3_’ ranging from 10 to 80 nt. Bases in black are for the 30 nucleotides flanking the SSR each side. **D,** Confirmation of mutation c.584T>C in additional weeping cultivars by direct sequencing of PCR amplicons. The target SNV at 584 in the sequence chromatogram is highlighted in black, which represents a heterozygous genotype T/C.

An interesting finding in the promoter sequences was a variable SSR region [(TA)_2-12_(TG)_0-25_(TA)_1-3_] located from −359 to −280 nt (upstream of the ATG site), and the three weeping cultivars had an identical SSR allele [(TA)_2_(TG)_0_(TA)_3_] that is 12-70 nt shorter than the other seven standard alleles (**Fig.3A, C, Supplemental Fig. S3**). Moreover, the three shortest SSR alleles from the weeping cultivars were also associated with the weeping allele *MdLAZY1A-W* (carrying CC_584_T) in coupling phase, explaining why marker Ch13-8547, which detects the SSR region, was reliable in genotyping the open-pollinated progenies of CW and RJ.

### Suppression of the standard allele *MdLAZY1A-S* expression changed branch growth direction to downward

To confirm the identification of *MdLAZY1A*, an RNAi construct targeting *MdLAZY1A* was constructed and transformed into apple cultivar Royal Gala (RG) of standard growth (*ww*) via *Agrobacterium*-mediated stable genetic transformation, generating six independent RNAi lines. Quantitative real-time PCR (qRT-PCR) analysis of their actively growing shoot apex tissues revealed that three of the six RNAi lines (1, 3 and 6) showed a significant reduction in the expression level of *MdLAZY1A* as expected (**Fig. 4A-D, G**). Measurements on 2-year-old trees indicated that the RNAi lines had significantly wider branch angles in relation to the vertical trunk (75.3°-79.7° vs 57.2°) and more spreading or downward branch orientations (93.8°-102.9° vs 44.5°) than the non-transgenic control plants **(Fig. 4A-F)**. These results demonstrated that suppression of allele *MdLAZY1A-S* leads to the downward growing branches, strongly supporting *MdLAZY1A* as the gene underlying *W* conferring weeping growth in apple.

**Figure 4.**
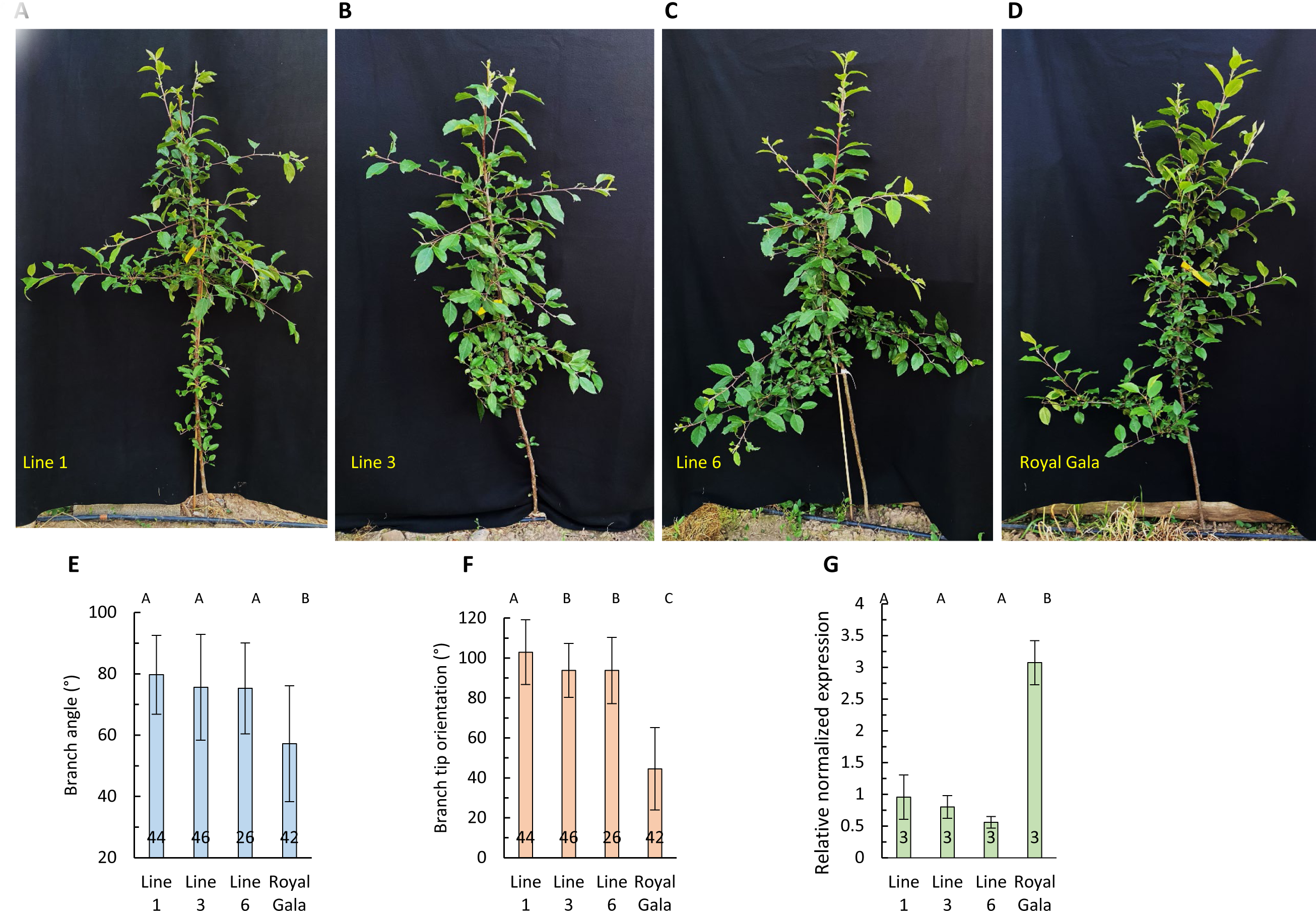
Suppression of standard allele *MdLAZY1A-S* in apple. **A-D,** Representative 2-year-old MdLAZY1A-RNAi transgenic (A-C) and non-transgenic Royal Gala (RG) (D) trees. **E-G,** Branch angle (E), branch tip orientation (F), and qRT-PCR analysis (G) of *MdLAZY1A-S* expression in apical shoot tips. The number of measurements or replicates (n) is given inside each chart. Error bars indicate standard deviation. Different letters denotes significant difference (p < 0.05) among lines or genotypes in Tukey’s HSD test.

### Genetic complementation of weeping allele *MdLAZY1A-W* led to weeping-like growth

To test if mutation c.584T>C (L195P) causes the weeping phenotype, the coding sequence of the weeping allele *MdLAZY1A-W* was isolated from CW and was cloned into an over-expression construct under control of the cauliflower mosaic virus 35S promoter. Genetic transformation by *Agrobacterium* was conducted similarly to transform the construct into RG, resulting in seven independent lines. Three lines (3, 4 and 5) with at least six clones per line were selected due to phenotype for further studies. At three months of age, leaf angles were measured for the transgenic lines and the non-transgenic RG plants. The data showed that the transgenic lines ranged from 104.6° to 111.3° (from the vertical trunk), significantly wider than that of non-transgenic RG (58.1°) **(Fig. 5A, C)**. At three years of age, the transgenic lines had spreading and downward oriented branches (**Fig. 5B**). Quantitatively, the branches from the three transgenic lines had significantly wider branch angles (76.4°-85.8°) and more spreading and downward branch orientations (84.5°-96.5°) than those from the non-transgenic RG (61.0° and 50.7°, correspondingly) (**Fig. 5D-E)**. qRT-PCR analysis of *MdLAZY1A* expression in apical shoot tips showed that all transgenic lines had significantly higher expression levels of the gene than RG (**Fig. 5F**), suggesting that over-expression of the weeping allele *MdLAZY1A-W* in RG led to spreading and downward leaves and branches, similar to the phenotype observed when the standard allele (*MdLAZY1A-S*) was knocked down by RNAi (**Fig. 4).**

**Figure 5.**
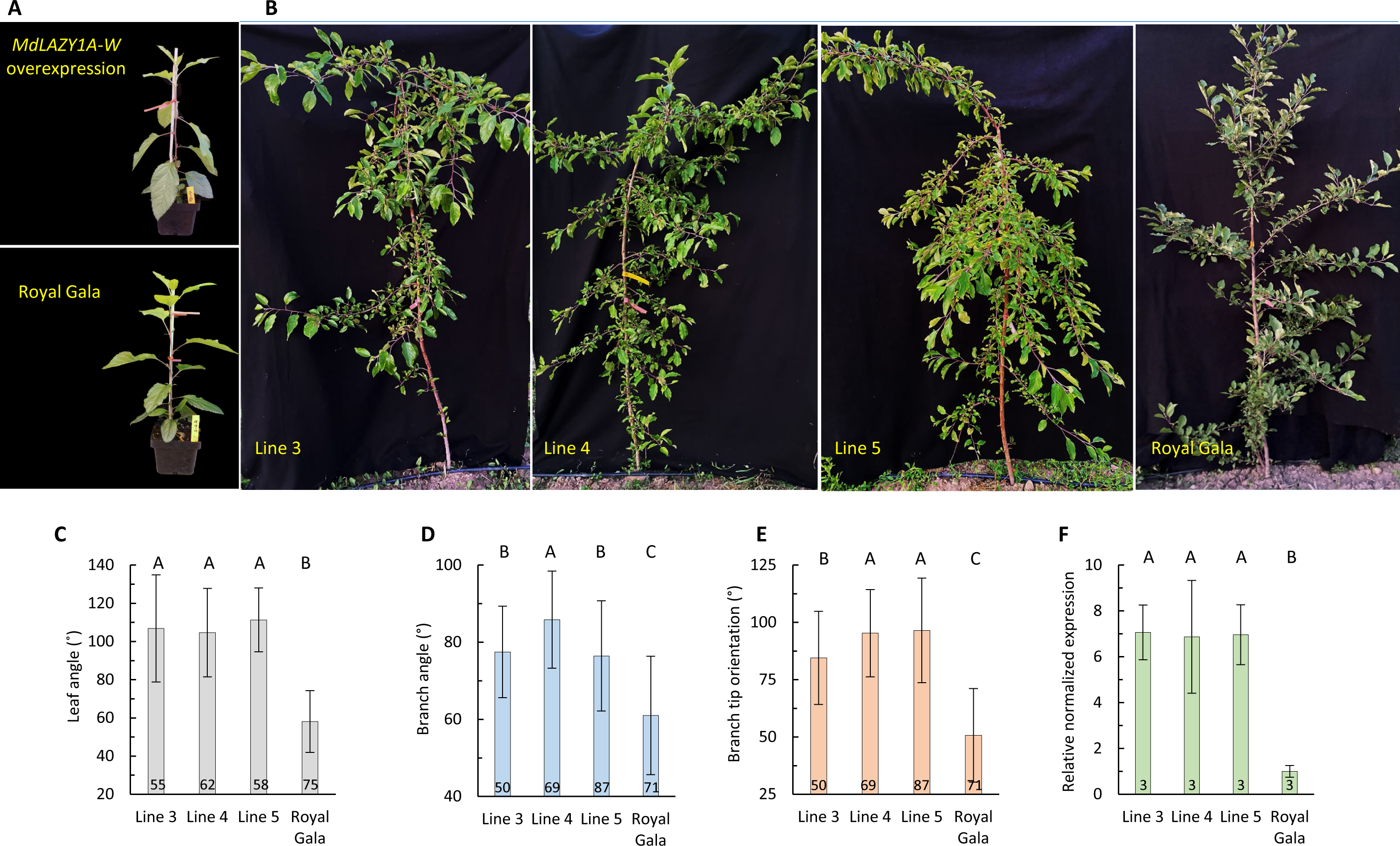
Overexpression of weeping allele *MdLAZY1A-W* in apple. **A-B,** Representative transgenic and non-transgenic Royal Gala (RG) trees at 3 months of age (A) and 3 years of age (B). **C,** Leaf angle of 3-month-old transgenic and non-transgenic RG trees. **D-E,** Branch angle (D) and branch tip orientation (E) of transgenic and non-transgenic RG trees at 3 years of age. **F,** qRT-PCR analysis of *MdLAZY1A* expression. The number of measurements or replicates (n) is given inside each chart. Error bars indicate standard deviation. Different letters denotes significant difference (p < 0.05) among the genotypes in Tukey’s HSD test.

To determine if over-expressing the standard allele would also confer a weeping-like phenotype, an over-expression construct of *MdLAZY1A-S* (isolated from CW) was constructed and transformed into a Royal Gala apple seedling selection (GL-3) reported to have high transformation efficiency (Dai et al., 2013), producing six independent transgenic lines. However, qRT-PCR analysis of their main shoot apex tissues indicated that only one line (L6) displayed the expected over expression of *MdLAZY1A*, while the other five lines exhibited either no significant change (L2 to L5), or under-expression (L1) (**Supplemental Fig. S5A-C, G**). The *MdLAZY1A* under expression line, which was presumably caused by co-suppression, had drooping leaves at a young stage (**Supplemental Fig. S5C**), similar to leaves observed in the *MdLAZY1A-W* overexpression lines (**Supplemental Fig. S5A**). However, the over expression line (L6) appeared to have more upward leaves than the non-transgenic control during early stages **(Supplemental Fig. S5A)**. When branches emerged, the over expression line (L6) had a significantly narrower branch angle (54.7°) and a more upward branch orientation (47.8°) than the non-transgenic control (branch angle 66.3°, branch orientation 58.1°), whereas the under expressed line (L1) had a significantly wider branch angle (87.4°) and a more spreading branch orientation (91.8°) than the non-transgenic control plants **(Supplemental Fig. S5D-F, H, I)**.

Overall, these results imply that the standard allele promotes upward branch growth, and its suppression may lead to downward branch growth, similar to what was observed in the RNAi lines (**Fig. 4)**, suggesting that the weeping phenotype in the *MdLAZY1A-W* over-expression lines was more likely caused by the presence of *MdLAZY1A-W* than by the elevated transcription of the allele. We concluded, therefore, that mutation L195P in MdLAZY1A is genetically causal for weeping growth in *Malus*.

### *MdLAZY1A-W* impairs shoot gravitropic response

Evaluation of the impact of *MdLAZY1A-W* on tree response to gravity was first conducted using the growing primary branches of both RG and the *MdLAZY1A-W* lines (Line 4 and Line 5) by forcing their tips to orient horizontally or 90° from their main truck (0°) for 4 days (**Supplemental Fig. S6**). Daily tracking of their branch tip orientations demonstrated that RG had a rapid upward trending in one day and continued progressively through day 4, similar to the gravitropic curvatures observed in *Populus* (Groover, 2016). In contrast, Line 4 and Line 5 exhibited a slow downward trending, suggesting a limited gravitropic response.

To investigate further the effect of *MdLAZY1A-W* on gravitropic response, a new set of the three *MdLAZY1A-W* transgenic lines (Lines 3-5) alongside the non-transgenic RG plants were raised and re-oriented horizontally when trees were 91.5±14.4 cm in height (**Fig. 6A**). Measuring their main shoot orientation from day 0 (D0) through D50 during the treatment uncovered that the transgenic lines lifted their growth direction slower and to a less degree than RG, although a statistically significant difference (in main shoot orientation) between RG and the transgenic lines was not detected at all time points for all lines (**Fig. 6A, B, F**). Three weeks after the trees were re-orientated horizontally, a few buds broke at the base area of the stems (close to roots) in RG and Lines 3-5 (2-3 days later than RG). By D47, the buds grew into 3-5 branches on each tree (**Fig. 6B**), prompting reversing the trees back to their original upward standing position at D51 (D0’) so that all branches were re-oriented horizontally (**Fig. 6C**). At D0’, the branches had similar branch length, angle, and orientation between RG and the *MdLAZY1A-W* overexpression lines, although the branches were shorter in Line 3 (**Fig. 6G-I**). During the following six weeks, the branches of Lines 3-5 showed significantly reduced upward growth than those of RG at all six time points measured (**Fig. 6G**). These data strongly implicated that the *MdLAZY1A-W* lines had an impaired gravitropic response in the elongation zone of both main shoots and branches.

**Figure 6.**
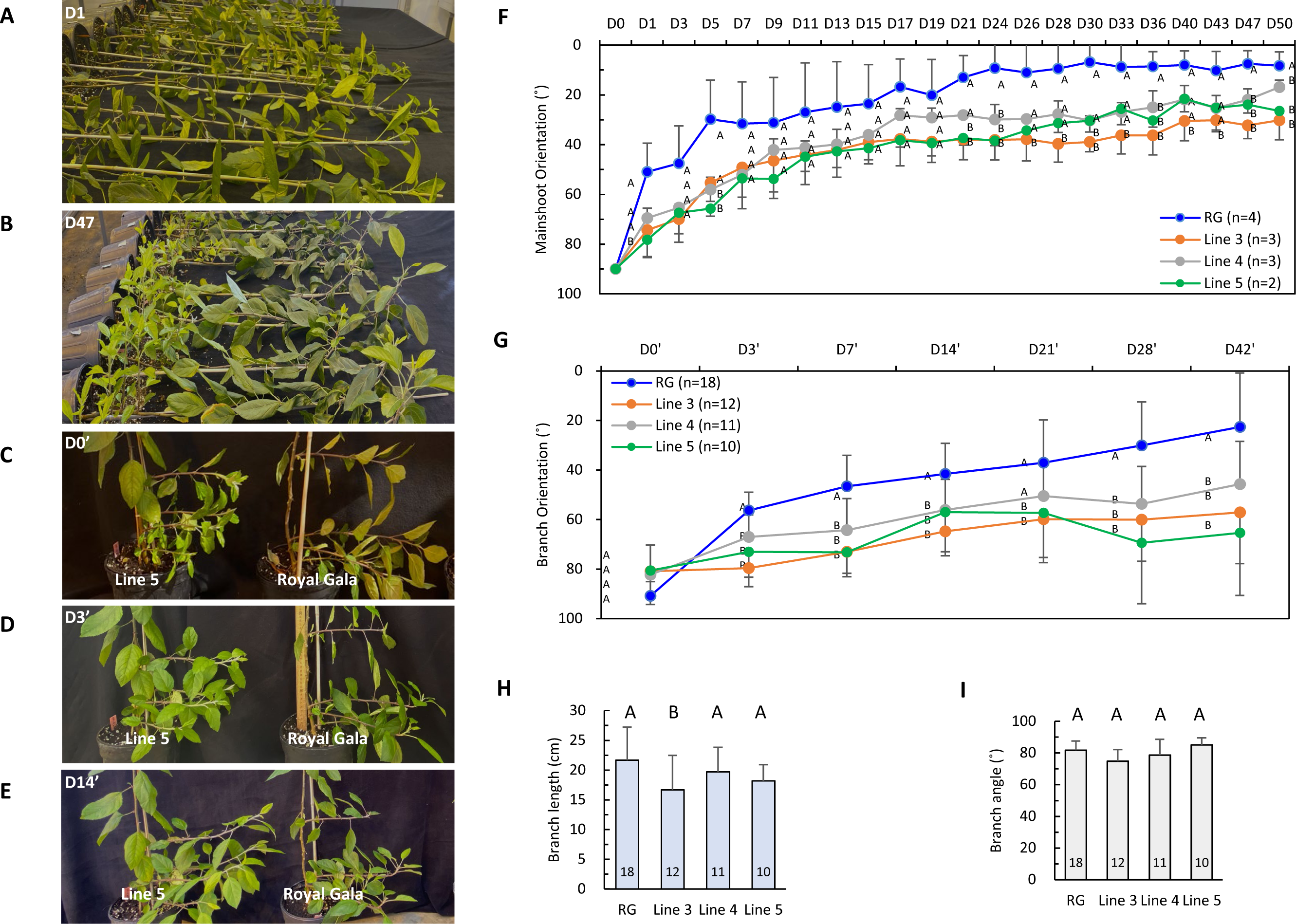
Evaluation of gravitropic response of the *MdLAZY1A-W* over-expression lines using young plants (5-month old). **A-B,** Photos of the transgenic lines and RG (non-transgenic control) trees at Day 1 (D1) (A) and D47 (B) after being re-oriented horizontally. Note the changes in main shoot tip orientation in A and B and emerging of new branches in B. **C-E,** Photos of the transgenic lines and RG at D0’ (day 0) (C), D3’ (D) and D14’ (E) after trees were reversed back to standing vertically. **F-G,** Measurements of main shoot tip orientation (F) and branch tip orientation (G). Significant difference in Tukey HSD test at each time point between Gala and Lines 3-5 were indicated by different letters. **H-I**, Branch length (H) and branch angle (I) measured at D0’. The number of branches (n) is given inside each chart.

### Mutation L195P resides within a conserved region of the predicted transmembrane domain in MdLAZY1A

To study the impact of mutation L195P, the protein sequence of MdLAZY1A-S was aligned with AtLAZY1 in *Arabidopsis* (Yoshihara et al., 2013) and OsLAZY1 in rice (Li et al., 2007; Yoshihara and Iino, 2007) (**Fig. 7**). The alignment showed that mutation L195P occurred in Region III, one of the five conserved regions in LAZY1-like proteins (Yoshihara et al., 2013), suggesting that the mutation might have important functional consequences. However, the amino acid residue corresponding to L195 is conserved in OsLAZY1 only, as it was substituted by amino acid phenylalanine (F) in AtLAZY1 (**Fig. 7**).

**Figure 7.**
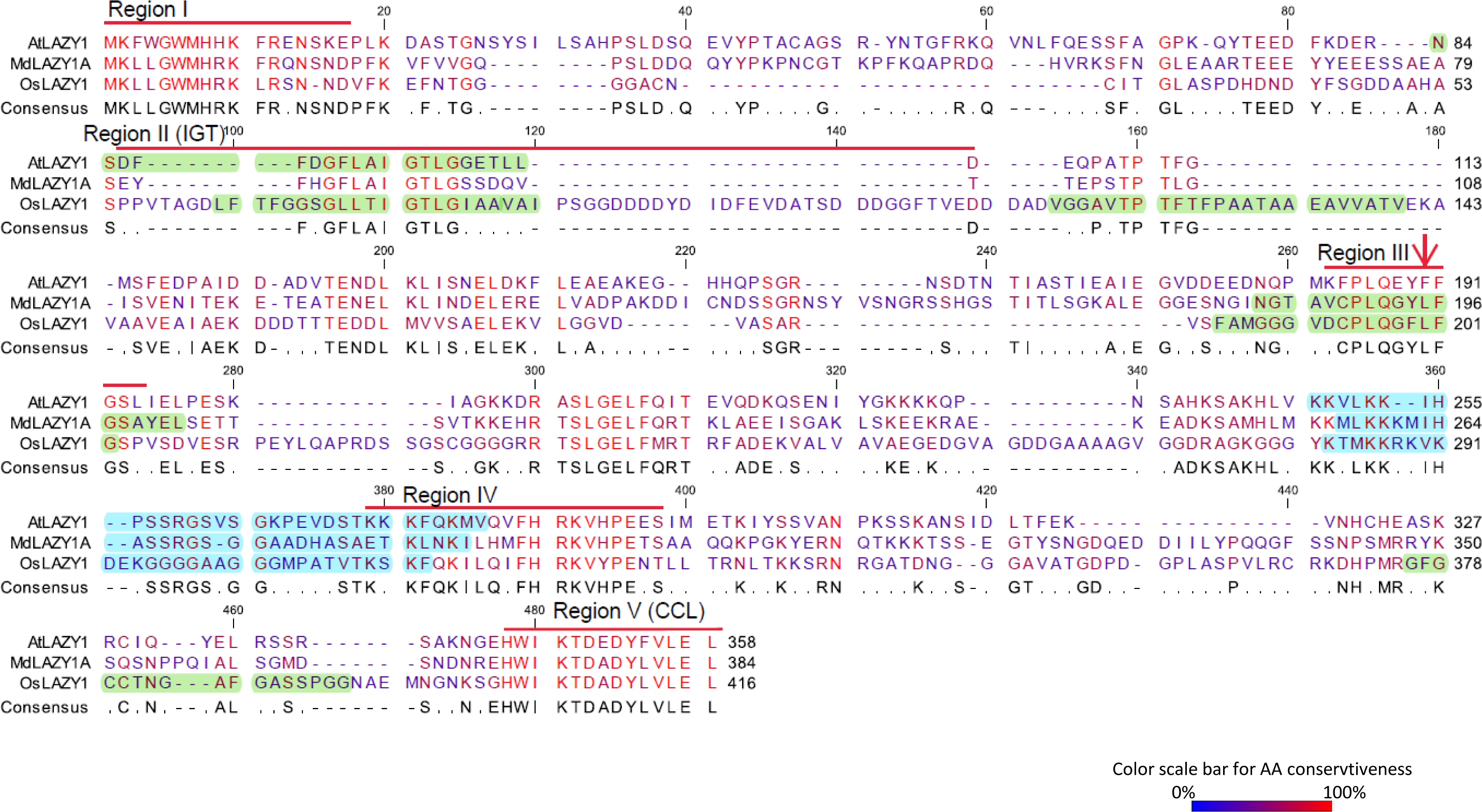
Alignment of MdLAZY1A protein sequence with those of AtLAZY1 (At5g14090) in *Arabidopsis* and OSLAZY1 (Os11g29840) in rice. The conservativeness of amino acids is shown in color as indicated by the color scale. The residues without consensus are indicated by dots. The five conserved regions as indicated by Yoshihara et al (2013) are shown with a thick up-line in red. Transmembrane domains predicted by TMpred are highlighted in green. Nuclear localization signals (NLS) predicted by cNLS Mapper were highlighted in blue. The L195P mutation in MdLAZY1A responsible for apple weeping growth is indicated with a red arrow.

To examine if and how residue L195 is conserved in other LAZY1 orthologs, a BLAST search against the NCBI databases of reference proteins (refseq-protein) was conducted using AtLAZY1, resulting in 420 significant hits (cutoff e=0.05). Notably, AtLAZY2 through AtLAZY6 of *Arabidopsis* (Taniguchi et al., 2017) did not meet the cutoff, indicating their orthologs were not among them. Of the 420 LAZY1-like proteins, 218 representatives were selected and analyzed in more detail, which covered 81 plant species in 75 genera (**Supplemental Table S7**). For *Malus*, four distinct LAZY1-like proteins were among the 218 selections, including MdLAZY1A and three others, named MdLAZY1B (MD16G1122900), MdLAZY1C (MD17G1002300) and MdLAZY1D (MD09G1008500) on chromosomes 16, 17 and 9, respectively (**Supplemental Fig. S7B**). Alignment of the 218 protein sequences confirmed the five conserved regions (I to V) as previously defined (Yoshihara et al., 2013) (**Supplemental Figs. S7 and S8**). Surprisingly, L195 was detected in 91.3% (199/218) of the LAZY1 orthologs, suggesting L195 is a highly conserved amino acid in Region III (**Supplemental Figs. S7A and S8, Supplemental Table S8**), supporting the significance of mutation L195P in MdLAZY1A. In the 19 LAZY1 proteins, where L195 was substituted with other amino acids or not present (**Supplemental Table S8**), 14 showed marked variations in Region III. The remaining five LAZY1 proteins include four with F195 (including AtLAZY1) and one with P195 in XP_018682759, a protein from *Musa acuminata* subsp. *malaccensis* (D’Hont et al., 2012). Given the relative high frequency of F195 (1.83% or 4/218), a hypothetical MdLAZY1A with substitution L195F was compared with MdLAZY1A-W using an online tool-Protein Variation Effect Analyzer (Provean) (Choi et al., 2012) to predict their functional effect. The results showed that L195P and L195F had PROVEAN scores −6.25 and −2.96, which are considered deleterious and tolerated (cutoff = 4.1, a recommended stringent score) substitutions, respectively, suggesting L195F might be a more tolerated substitution than L195P.

Both MdLAZY1A-S and MdLAZY1A-W seemed to contain a transmembrane domain (TMD) (aa 184 to 202) (**Fig. 7, Supplemental Table S9**) and a bipartite nuclear localization signal (NLS) (aa 256-286) (**Fig. 7)** based on the TMD prediction program TMpred (Hofmann and Stoffel, 1993) and an NLS identification program cNSL Mapper (Kosugi et al., 2009). Although their associated TMpred scores were below the default significance cutoff of 500, the score of 488 for MdLAZY1A-S was close to the threshold while the score of 194 for MdLAZY1A-W was considerably lower (**Supplemental Table S9**). Interestingly, the predicted TMD encompassed Region III (**Fig. 7**) where mutation L195P occurred, implicating that the mutation is likely even more consequential.

To experimentally determine the MdLAZY1A cellular localization in *planta*, the *MdLAZY1A-S* and *MdLAZY1A-W* coding sequences were C-terminal tagged with green florescence protein (GFP) initially, and were transiently expressed in the leaves of *Nicotiana benthamiana* through *Agrobacterium*-mediated infiltration. Confocal microscopy revealed that both MdLAZY1A-S::GFP and MdLAZY1A-W::GFP showed a signal from the plasma membrane (PM) (**Supplemental Fig. S9A, B**) based on their co-localization with an mCherry tagged PM marker (PIP2A) (Nelson et al., 2007). A nuclear signal was also sporadically detected for both MdLAZY1A fusions (**Supplemental Fig. S9C-D**). Similar PM localization was observed under confocal microscope (**Fig. 8A, B**) when the MdLAZY1A was tagged at the N-terminal with a yellow florescence protein (YFP::MdLAZY1A-S and YFP::MdLAZY1A-W). Detailed analysis of images indicated that the YFP::MdLAZY1A chimeras localized to the periphery of the cells. Since the vacuole occupies a large volume in the leaf epidermal cell of *N. benthamiana*, it is challenging to distinguish plasma membrane from tonoplast. To address this, YFP::MdLAZY1A was co-expressed alongside the mCherry tagged PM (PIP2A) and tonoplast (γ-TIP) markers, independently (Nelson et al., 2007; Ivanov and Harrison, 2014), revealing YFP::MdLAZY1A predominant co-localization with the PM marker, as the strand-like structures associated with the invagination of vacuoles that were detectable by the tonoplast marker could not be detected by the PM marker (**Fig. 8A**). More importantly, the YFP::MdLAZY1A chimeras signal also co-localized with the PM marker protein in *Arabidopsis* Columbia protoplasts transiently co-transformed with the chimeras and the PM marker (**Fig. 8B**), thereby confirming that the PM localization of proteins MdLAZY1A-S and MdLAZY1A-W, although a conclusive difference in signal intensity between the two proteins could not be drawn.

**Figure 8.**
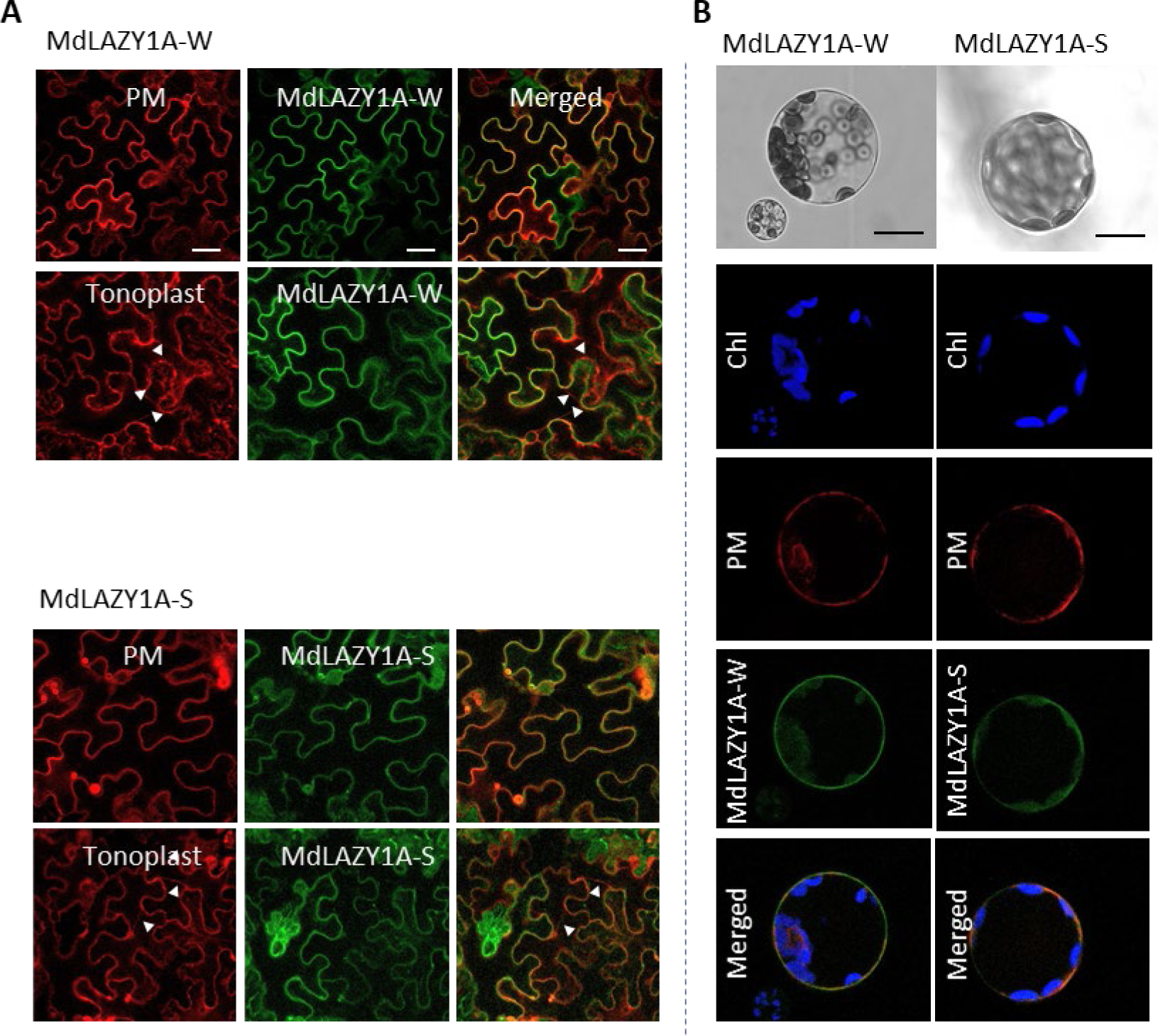
Confocal microscopy of the subcellular localization of MdLAZY1A-W and MdLAZY1A-S *in planta*. The YFP::MdLAZY1A-W and YFP::MdLAZY1A-S chimeras were transiently expressed in *Nicotiana benthamiana* epidermal cells (A) and *Arabidopsis* protoplasts (B). YFP is false-colored green throughout the images in A and B. **A,** The YFP::MdLAZY1A-W and YFP::MdLAZY1A-S chimeras transiently co-expressed with either the mcCherry tagged plasma membrane PM (PIP2A) or mcCherry tagged tonoplast (γ-TIP) membrane markers. The left panel shows the mCherry signal associated with the membrane markers, the middle panel shows the YFP signal (false-colored green) associated with the MdLAZY1A proteins, and the right panel shows the merged channels. The white triangles highlight the strand-like structures associated with the invagination of vacuoles. The scale bar represents 40 μm. **B,** The YFP::MdLAZY1A-W and YFP::MdLAZY1A-S chimeras (false-colored green) co-localized with the Arabidopsis PM marker protein (PIP2A-mCherry) when co-expressed in *Arabidopsis* mesophyll protoplasts. Chl shows the signal associated with chloroplast autofluorescence and is false-colored blue. Scale bars represent 20 μm.

### MdLAZY1A is the only LAZY1 paralog abundantly expressed in apple shoot

Phylogenetic analysis of the 218 LAZY1-like proteins suggested that they could be grouped into eight clades (I through VIII), and the largest clade (II) could be divided further into three sub-clades (II-A, II-B and II-C) (**Supplemental Fig. S7B**). Members in clade III were mostly from monocots, including OsLAZY1 from rice (Li et al., 2007; Yoshihara and Iino, 2007) and ZmLAZY1 from maize (Dong et al., 2013). Interestingly, the four apple LAZY1 proteins were members of two different clades: MdLAZY1A and MdLAZY1B were in clade VI, whereas MdLAZY1C and MdLAZY1D were in clade II-B alongside AtLAZY1, suggesting MdLAZY1A and MdLAZY1B are less related to AtLAZY1 than MdLAZY1C and MdLAZY1D. To see why MdLAZY1A was determined to be instrumental in weeping growth while the others were not, a search for non-synonymous SNVs specific to weeping in genes *MdLAZY1B, C,* and *D* was attempted in the weeping and standard pooled genome sequences used to map the weeping phenotype previously (Dougherty et al., 2018). However, such non-synonymous SNVs were not found. Moreover, based on the RPKM values in RNA-seq analysis, *MdLAZY1A* (15.8-20.0) was expressed abundantly while *MdLAZY1B* (0.5-1.2), *MdLAZY1C* (0.7-0.8) and *MdLAZY1D* (0-0.2) had barely detectable transcripts (**Fig. 9A, Supplemental Table S3**). Furthermore, comparing the 2-kb promoter region upstream of the ATG site in *MdLAZY1A* with the corresponding regions in *MdLAZY1B, C,* and *D* revealed no significant similarities among them, suggesting the promoter of *MdLAZY1A* is unique, and likely responsible for its strong expression in shoots. To determine how *MdLAZY1A* was expressed in organs of apple (a weeping progeny of CW), a qRT-PCR analysis was conducted, revealing that the gene expression was comparable in leaf, shoot and flower, but was 6.1-8.5-fold lower in the roots (**Fig. 9B**), suggesting it was primarily expressed in the above-ground organs of apple.

**Figure 9.**
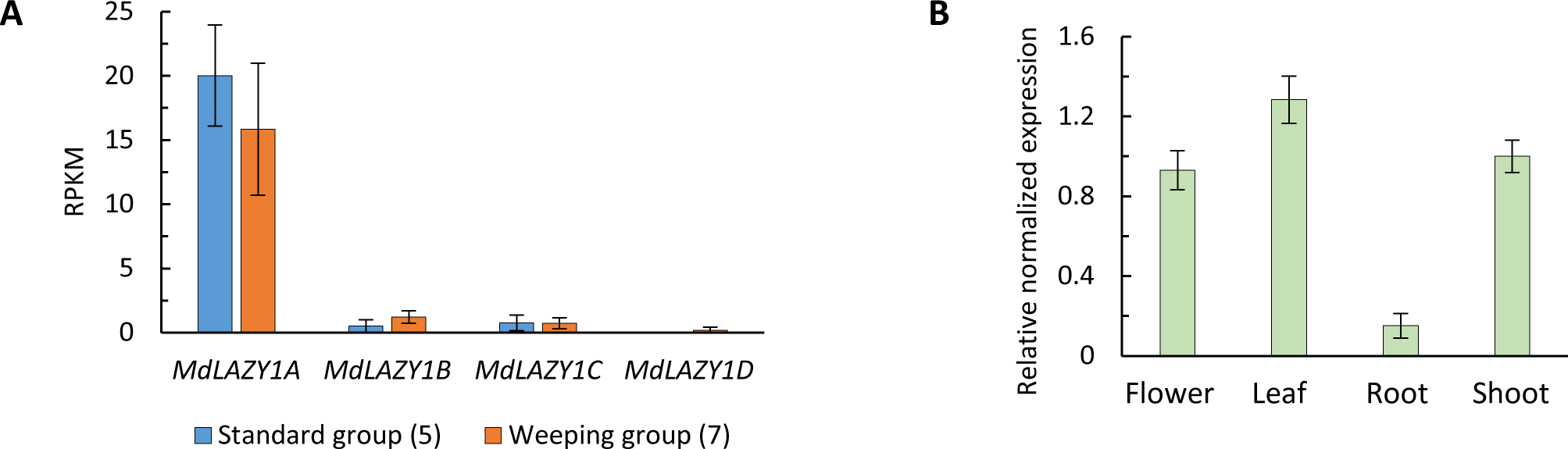
*MdLAZY1A* expression assays. **A,** Expression of *MdLAZY1A* and its paralogs *MdLAZY1B, C* and *D* in a shoot apex tissues as determined by RNA-seq. *MdLAZY1A=MD13G1122400; MdLAZY1B=MD16G1122900; MdLAZY1C=MD17G1002300; and MdLAZY1D=MD09G1008500*. **B,** Expression of *MdLAZY1A* in different organs in apple (a weeping progeny of CW). The root tissue was taken from its seedling roots.

## Discussion

### L195P is likely a dominant-negative mutation of MdLAZY1A

Weeping-like phenotypes in plants mostly result from a recessive loss-of-function mutation, such as *lazy1* in rice (Li et al., 2007; Yoshihara and Iino, 2007) and *weep* in peach (Hollender et al., 2018). In the case of *WEEP*, which encodes a conserved protein containing a sterile alpha motif (SAM) domain, the loss-of-function mutation was caused by a 1.8-kb deletion abolishing the 5’ end of the gene (Hollender et al., 2018). For *LAZY1*-like genes, various recessive loss-of-function mutations were reported, including short deletions induced loss of LAZY1 C-terminus (Li et al., 2007; Yoshihara and Iino, 2007) and a large deletion prompted *LAZY1*-null (Yoshihara and Iino, 2007) in rice, a single nucleotide mutation (at the first splicing junction) generated loss of LAZY1 N-terminus (Dong et al., 2013) and short InDels caused frame shift and premature stop codons (Dong et al., 2013; Zhang et al., 2014) in maize, and T-DNA insertion induced truncation of LAZY1 N-terminus in *Arabidopsis* (Yoshihara et al., 2013). In silver birch (*Betula pendula*), an in-frame point mutation in the *BpLAZY1* gene converting serine-44 into a stop codon was found in the weeping birch ‘Youngii’, prompting speculation that weeping growth in silver birch resulted from a loss-of-function mutation in *LAZY1* (Salojärvi et al., 2017).

In contrast, the weeping phenotype in *Malus* is dominant over the standard growth phenotype, which is rare in plants. One exception was in Japanese chestnut (*Castanea crenata*) where a dominant weeping allele was mapped, although a recessive weeping allele was more prevalent in the germplasm (Terakami et al., 2021). Since the dominant weeping phenotype in *Malus* is largely a result of a single amino acid substitution L195P in MdLAZY1A-W, the mutation was likely to be exceedingly detrimental to the weeping allele function so that it even prevented the standard allele MdLAZY1A-S from functioning in heterozygous plants. Indeed, the predicted Provean scores (Choi et al., 2012) for amino acid substitution L195P in MdLAZY1A was deleterious (−6.25), while the score predicted for the hypothetical substitution L195F (present in the wild type allele of *Arabidopsis*) in MdLAZY1A was tolerated (−2.96), supporting the notion that L195P is a detrimental mutation in MdLAZY1A.

Intra-locus dominant mutations are categorized into three major groups: (1) dominant negative mutations (DNMs), (2) haploinsufficiency (HI)/reduced gene dosage, expression or protein activity, and (3) increased gene/gene-product dosage or activity (gain-of-function) (Veitia et al., 2018). DNMs have been documented in other plant species, such as the mutation *OsKAT2* encoding an inward rectifying shaker-like potassium channel protein in rice (Moon et al., 2017), *Nonphototropic seedling1* (*Nps1*) encoding a PHOT1 like phototropin in tomato (Sharma et al., 2014; Kilambi et al., 2021) and *ML14257* encoding an actin in monkeyflower (*Mimulus lewisii)* (Ding et al., 2017). In *Arabidopsis*, a survey of dominant phenotype mutations induced by chemical ethyl methanesulfonate (EMS) in 92 genes concluded that these mutations were caused mostly (73.9% or 68/92) by missense mutations that do not increase transcription, but change protein function or have a dominant-negative effect on protein-protein interactions (Meinke, 2013). Based on the mutation characteristics, L195P is considered a dominant negative mutation, as the protein of the mutation allele appears to disrupt the function of the standard allele when they co-exist in cells of heterozygous individuals of weeping growth.

### Implications of Mutation L195P in MdLAZY1A-mediated gravitropism signaling

The MdLAZY1A proteins were localized to the plasma membrane and nucleus in plant cells (**Fig. 8, Supplemental Fig. S9**), consistent with the subcellular localization of LAZY1 proteins reported in rice (Li et al., 2007), *Arabidopsis* (Yoshihara et al., 2013), and maize (Dong et al., 2013). Given that the NLS in OsLAZY1 has been confirmed in rice (Li et al., 2007; Li et al., 2019), the predicted NLS sequence in MdLAZY1A, which was aligned across MdLAZY1A, AtLAZY1 and OsLAZY1 (**Fig. 7**), could be considered responsible for its nuclear location. However, mutation L195P is not present in the NLS region of MdLAZY1A, suggesting that the nucleus would unlikely be a site of action for MdLAZY1A-W. Nevertheless, the importance of nucleus localization of LAZY1 proteins in shoot gravitropism is reportedly varied depending on species, ranging from not being required in *Arabidopsis* (Yoshihara et al., 2013; Yoshihara and Spalding, 2020) to potentially important in maize (Dong et al., 2013) and essential in rice (Li et al., 2019).

Proteins MdLAZY1A-W and MdLAZY1A-S were predicted to have an identical TMD position (aa 184-202) in the conserved Region III by TMpred (Hofmann and Stoffel, 1993) with scores of 488 and 194, respectively (**Fig. 7, Supplemental Table S9**). Although both scores were below 500, the significance cutoff (Hofmann and Stoffel, 1993), the TMD predicted in MdLAZY1A-S is close to be significant while that of MdLAZY1A-W is not, suggesting (again) a considerable adverse effect of the L195P mutation on the predicted transmembrane helix. This is congruent with the negative impact of the L to P substitution on TMDs reported in other species, such as wheat (Lillemo and Morris, 2000), barley (Civáň and Brown, 2017), and humans (Molnár et al., 2016), although the L195P mutation influence on MdLAZY1A’s association with PM remains unclear (**Fig. 8, Supplemental Fig. S9**).

OsLAZY1 was predicted with four TMDs in present study, designated TMD1 (aa 62-83, in Region II), TMD2 (aa 117-140), TMD3 (aa 186-202 in Region III) and TMD4 (aa 376-392), of which TMDs 1, 3 and 4 were associated with significant scores (523-1745), while AtLAZY1 was predicted with a single TMD (aa 84-103 in Region II) of a low and insignificant score of 37 (**Fig. 7, Supplemental Table S9**). This indicated that both TMD1 in OsLAZY1 and the single TMD in AtLAZY1 were predicted to be in Region II or the IGT region (Dardick et al., 2013), whereas the TMD3 in OsLAZY1 and the single TMD in MdLAZY1A were in Region III. Since the prediction of TDM1 and TMD4 in OsLAZY1 were largely confirmed by the observation that deletion of the N-terminal region (aa 1-100) or C-terminal region (aa-321-416) led to a loss of its plasma membrane localization (Li et al., 2007; Li et al., 2019), the predicted TMD3 in OsLAZY1 and its counterpart in MdLAZY1A in Region III would likely be plausible.

In a study systematically substituting two amino acids in all five conserved regions in AtLAZY1 (Yoshihara and Spalding, 2020), the resultant variant (P185A/ L186A) in Region III showed typical wild-type subcellular localization, and also rescued the branch angle defect in *atlazy1*, indicating that such substitution in Region III did not have significant effect on gravitropic response in *Arabidopsis*, while similar substitutions in Regions I, II and V had a dramatic consequence (Yoshihara and Spalding, 2020). These findings in *Arabidopsis* contradict the observation that mutation L195P in Region III causes weeping growth in apple. A few factors might have contributed to such discrepancy. First, mutation P185A/ L186A in AtLAZY1A is equivalent to P190A/ L191A in MdLAZY1A, different from mutation L195P (**Fig. 7**). Second, L195 in MdLAZY1A corresponds to F190 in AtLAZY1, which is considered a mutation from L190 already (**Fig. 7, Supplemental Fig. S8**). Third, the amino acid sequences flanking Region III in AtLAZY1 are so different from those in MdLAZY1A and OsLAZY1, as a TMD was predicted in Region III in both MdLAZY1A and OsLAZY1, while not in AtLAZY1. Lastly, apple, as a plant, diverges drastically from Arabidopsis. A dedicated study is needed to understand the seemingly contradictory findings.

Disruption of protein homo-dimer and competition for substrate were considered the common molecular mechanisms underlying DNMs (Wilkie, 1994; Veitia et al., 2018). Since 65% of membrane proteins were estimated to be obligate oligomers (Forrest, 2015), it is possible that MdLAZY1A might function as a homo-dimer in cells and the L195P mutation might result in non-functional complexes. As such, the mutation would affect the dimer configuration of the MdLAZY1A protein complex on the plasma membrane, thereby its interactions with other gravitropism signaling components, such as the RLD proteins that regulate the polar re-localization of PIN proteins (Furutani et al., 2020), the AGCVIII protein kinase-like PKC that may phosphorylate PINs and other polar auxin transporters (PATs) (Dong et al., 2013), and the BRX proteins critical in gravitropism in plants (Li et al., 2019; Che et al., 2023). Future experiments are needed to determine the mechanism by which MdLAZY1A-W impairs gravitropic responses in apple cells.

### The L195P mutation site may serve as a target of base editing for tree architecture improvement in apple and beyond

Although there are at least 7,500 apple varieties named (Elzebroek and Wind, 2008), apple production worldwide has been dominated by approximately 30 apple cultivars, such as Gala, Fuji among others. These cultivars have upward branches, requiring extensive training, such as pruning and branch bending to optimize tree canopy for maximum productivity and profitability (Ferree and Schupp, 2003; Musacchi and Greene, 2017). Downward branch bending has been practiced for centuries in orchards due to the beneficial effect of increasing flowering and reducing vegetative growth (Wareing and Nasr, 1958). However, such branch bending in commercial orchards is labor-intensive and costly. To reduce production costs, it is desirable to genetically alter the widely grown apple cultivars from growing branches upward to spreading or downward, but not to change their identity that has been critical to consumer preferences. Ideally, such improvement could be accomplished without integration of any exogenous DNA to be potentially exempted from regulations (Menz et al., 2020). With the advances in clustered regularly interspaced short palindromic repeats (CRISPR)-associated-protein (CAS) genome editing technologies, such as base editing (Gaudelli et al., 2017), prime editing (Anzalone et al., 2019) and DNA-free gene editing (Tsanova et al., 2021), the finding of the L195P mutation opens up a plausible path for the desired improvement. It is known that CRISPR-CAS based gene editing in plants preferentially generates heterozygous mutations in T_0_ (Jin et al., 2021; Lu et al., 2021). Successful DNA-free base editing for mutation c.584T>C would create a dominant negative allele of *MdLAZY1A* in T_0_ apple plants, which are expected to grow downward branches, thereby rapidly achieving the improvement of existing cultivars. Although technical challenges likely remain in exploring this proposed utility of mutation L195P, it is encouraging that successful base editing has been demonstrated in apple (Malabarba et al., 2021) and a base-edited *LAZY1* has been reported in rice (Li et al., 2021).

In the 218 LAZY1-like proteins aligned, the only other LAZY1 protein that contains a mutation equivalent to L195P was accession XP_018682759 (**Supplemental Figs. S7 and S8, Supplemental Table S7**), which is one of the three LAZY1 proteins encoded in the genome of wild banana (*Musa acuminata* subsp. *malaccensis*) (D’Hont et al., 2012). Coincidentally, banana inflorescence stems are known for their downward bending shortly after emergence due to their changes in gravitropic response from negative to positive (RAM et al., 1962). Given the low frequency of the specific mutation in LAZY1-like proteins, we speculate that L195P might have been responsible for such developmentally regulated reversal to gravitropic response in banana. If this were confirmed, the L195P mutation likely would have a similar effect on gravitropism in other plant species beyond apple, broadening its applicable range in genetic improvement of crop plant architecture, particularly in fruit and nut tree species.

## Conclusions

*MdLAZY1A* is the genetic determinant underlying the *W* locus controlling weeping growth in *Malus*. A single nucleotide mutation of c.584T>C leading to amino acid substitution L195P in the conserved Region III of MdLAZY1A is sufficient to disrupt the function of MdLAZY1A in both alleles in heterozygous plants and alter apple tree growth from standard to weeping-like. Amino acid L195 and Region III in MdLAZY1A play a crucial role in MdLAZY1A-mediated gravitropism and overall tree architecture in apple. This study sheds light on woody plant gravitropism and offers a critical entry point for genetic improvement of tree growth habit in apple.

## Materials and methods

### Plant materials, phenotyping and fine mapping of the *Weeping* (*W*) locus

A set of five populations segregating for the weeping phenotype in 1246 seedling trees were used for fine mapping of the *W* locus on chromosome 13 (**Table S1**). They included three F_1_ populations reported previously (Dougherty et al., 2018), which were developed from crosses of Cheal’s Weeping (CW) (a weeping crabapple cultivar) × Evereste, NY-051 × Louisa (a weeping crabapple cultivar) and NY-011 × NY-100 (a weeping selection derived from CW) that comprised 38, 140 and 39 seedlings, respectively. However, the number of F_1_ progenies in population CW × Evereste was expanded from 38 to 98 in this study. The remaining two were developed from open-pollinated progenies of seed parents CW and Red Jade (RJ), consisting of 738 and 231 seedlings, respectively (**Supplemental Table S1**). The seedling trees were phenotyped as weeping (W), weeping-like (WL), standard (S) or standard-like (SL) **(Supplemental Fig. S10)** as described previously (Dougherty et al., 2018). Weeping trees had almost all branches growing downwards, while weeping-like trees had a majority of branches growing downward. Standard trees had upward growth of most branches and standard-like trees had a majority of upward branches **(Supplemental Fig. S10).** Additionally, eight diverse weeping crabapple cultivars were also used, including two (Cascade and Oekonomierat Echter-meyer) from the USDA national *Malus* germplasm repository, Geneva, New York, and six (Molten Lava, Ludwick, A-23, Beverly NSY, Sinai Fire and Ann E. Manback weeper) from the Newman Arboretum, Cornell Botanic Gardens, Ithaca, New York.

DNA was extracted from leaf tissue using a cetyl trimethyl ammonium bromide (CTAB) method (Doyle and Doyle, 1987), and assays of SSR markers were conducted as described previously (Wang et al., 2012). High resolution melting (HRM) markers were developed and analyzed according to a procedure outlined earlier (Ban and Xu, 2020), which target the single nucleotide polymorphisms (SNPs) within the *W* region of 982-kb (Dougherty et al., 2018) **(Supplemental Table S2).**

For detection of the weeping allele *W*, two markers were used across all populations, i.e., SSR marker Ch13-8547 located in the *MdLAZY1A* promoter region and HRM marker H_9031 that detects the weeping causal mutation c.584T>C in *MdLAZY1A* (**Supplemental Table S2**). However, due to distinct genetic backgrounds in CW, RJ and Louisa, genotypic screening for informative recombinants in the *W* region was conducted with different sets of flanking markers. They included HRM markers H_8694 and H_9574 for the progeny of CW, SSR markers Ch13-8181 and Ch13-9530 for the progeny of Louisa, and H_8735 and Ch13-8547 for the RJ related progenies (**Fig. 2, Supplemental Table S2**). To support the identification of mutation c.584T>C, genome sequences of 28 diverse *Malus* accessions of standard growth were downloaded from NCBI-SRA databases (Duan et al., 2017).

### Determination of the genomic and cDNA sequences of *MdLAZY1A* alleles

The *MdLAZY1A* genomic DNA sequences of three weeping cultivars CW, RJ and Louisa and two standard apples Golden Delicious and NY-051 were determined by PCR amplification using primers LAZY1F/R_S (**Supplemental Table S2**). The amplicons were cloned independently into vector pJET1.2/blunt (ThermoFisher Scientific, Waltham, MA) and Sanger sequenced at Cornell University Biotechnology Resource Center, Ithaca, New York.

To determine the cDNA sequences of *MdLAZY1A* alleles, actively growing shoot apex tissues were sampled from CW and were flash-frozen in liquid nitrogen and stored at −80°C until use. RNA was isolated from one gram of ground tissues as previously described (Meisel et al., 2005) with slight modifications. Briefly, ground tissues were mixed with CTAB buffer and incubated for 30 minutes at 65°C, and then centrifuged. Supernatant was mixed with equal amounts chloroform and centrifuged again. RNA was precipitated from the resultant supernatant in lithium chloride at −20°C for 2-hour and re-suspended in 87.5 µl of water. The RNA samples were treated with DNase I (amplification grade, Invitrogen, Carlsbad, CA) and cleaned with RNeasy MinElute Clean up Kit (Qiagen, Hilden, Germany). RNA concentrations were determined using NanoDrop 1000 (Thermo Fisher Scientific) and RNA integrity was assayed on 1% agarose gel. One microgram of total RNA was used in reverse transcription reactions using the Superscript III RT (Invitrogen, Carlsbad, CA, USA) to obtain the first strain cDNA. The coding sequences of alleles *MdLAZY1A-S* and *MdLAZY1A-W* with or without the stop codon were PCR-amplified from CW cDNA using primers LAZY1F/R_S and LAZY1F/R_NS, respectively **(Supplemental Table S2)**. The PCR products were cloned into Gateway entry vector pCR8/GW/TOPO (Invitrogen, Carlsbad, CA, USA) and were sequence-confirmed.

### RNA sequencing

RNA-seq analysis was conducted with actively growing apical shoot tips sampled from 12 individuals, including four (2-weeping, 2-standard) from the CW family and eight (5-weeping, 3-standard) from the RJ family (**Supplemental Tables S3 and S4**). The shoot tissues were flash frozen and stored at −80°C until use. The total RNA was extracted, purified and quantified as described above. mRNA was isolated with NEBNext Poly(A) mRNA Magnetic Isolation Module and used to construct RNA-seq libraries with NEBNext Ultra Directional RNA Library Prep Kit for Illumina (New England Biolabs, Ipswich, MA) according to manufacturer’s protocol with slight modifications. 10 mg of the total RNA was used for mRNA isolation and one half of the isolated mRNA was used for library preparation. Libraries were multiplexed in equal amount for single-end 76-base sequencing by NextSeq 500 (Illumina, San Diego, CA) at the Cornell University Biotechnology Resource Center (Ithaca, NY). RNA-seq data analyses, including reads clean-up, reads mapping against the apple reference genome (Daccord et al., 2017) and quantification of gene expression between the weeping and standard progenies was conducted using the CLC Genomic workbench software (Qiagen, Beverly, MA) following a procedure described previously (Bai et al., 2015). Single nucleotide variants (SNVs) calling and synonymous and non-synonymous SNV identification in the two families were also conducted using the CLC Genomic workbench software (**Fig. 2**).

### Over- and under-expression analysis of *MdLAZY1A* in apple

For over-expression, the CDS clones (with the stop codon in vector pCR8/GW/TOPO) of alleles *MdLAZY1A*-*W* and *MdLAZY1A*-*S* were transformed into the plant overexpression vector pGWB412 (Nakagawa et al., 2007) by LR reaction and were sequence-confirmed, generating constructs MdLAZY1A-W:pGWB412 and MdLAZY1A-S:pGWB412. For under-expression, an *MdLAZY1A* RNAi construct was created following previously described guidelines (Helliwell and Waterhouse, 2003). Briefly, a 459 bp DNA fragment in the third exon of *MdLAZY1A* covering the CDS from 485 through 943 bp was PCR-amplified using primers with a flanking attB site (attB1-LAZY1-F and attB2-LAZY1-R, **Supplemental Table S2**), cloned into vector pCR8/GW/TOPO and sequence-confirmed. Clones of correct sequence were transformed into pHELLSGATE2 RNAi vector (Helliwell and Waterhouse, 2003) by BP reaction, creating the MdLAZY1A-RNAi construct. The MdLAZY1A-RNAi construct was restricted with enzymes *Xba* I and *Xho* I (New England Biolabs, Ipswich, MA) for two hours at 37°C and run on a 1.5% agarose gel to confirm insert size. Sequencing confirmation of the constructs was completed with primers P27-5 and P27-3 (**Supplemental Table S2**).

Apple cultivars Royal Gala (RG) and GL-3 [a RG seedling selection of high transformation efficiency (Dai et al., 2013)] with standard growth were used for stable genetic transformation. The apple transformation procedure described previously (Borejsza-Wysocka et al., 1999) was followed. PCR confirmation of the presence of the over-expression T-DNA in transgenic lines was conducted with primers pGWB35SF1 and LAZY1R-S (**Supplemental Table S2**), while that of the RNAi T-DNA by primers pGWB35SF1 and XM1688Ex3-R (**Supplemental Table S2).** Their construct plasmids and the non-transgenic RG and GL-3 were used as positive and negative controls, respectively.

For transgene expression analysis, qRT-PCR was performed using iTaq Universal SYBR Green Supermix (BioRad, Hercules, CA) on a CFX96 Touch Real-Time PCR Detection System (BioRad) following manufacturer’s protocol. The RNA samples were prepared from actively growing apical shoot tips of transgenic lines and non-transgenic controls with three biological replicates, and an apple actin encoding gene (EB136338) was used as a reference gene **(Supplemental Table S2).** The expression levels of the *MdLAZY1A* gene were quantified with the normalized expression (ΔΔCq) of the reference gene using the Bio-Rad CFX Maestro software.

Quantitative measurements for leaf angle, branch angle, and main shoot and branch tip orientation were recorded for the transgenic lines and non-transgenic controls, which were all grown on their own seedling roots from tissue culture. For each construct, measurements were taken normally from three transgenic lines with at least three clones per line. All measurements assumed that 0° is straight up, 90° is horizontal and 180° is straight down (**Supplemental Fig. S11**).

### Evaluation of gravitropic response

The transgenic lines over-expressing the weeping allele *MdLAZY1A-W* and the non-transgenic RG plants were evaluated for their response to gravity in two different settings. The first was the actively growing primary branches on 3-year-old trees of Line 4, Line 5 and RG. The trees were grown in pots in open environment during a regular growth season. For each genotype or line, 14 to 24 branches were selected from five trees and were forced to the horizontal position or 90° from the vertical main truck (0°) for four days. During this period, their branch tip orientations (°) were measured daily (**Supplemental Fig. S6**).

The second was a set of 12 young (5 months) trees, including three trees from Lines 3 and 4 each, two from Line-5, and four from RG. The trees, which were grown in greenhouse throughout the experiment, were placed sideways on the bench so that the main shoot tips were adjusted to 90° from a vertical axis (**Fig. 6A)**. The main shoot orientation (°) was measured every 1-3 days for 50 days in total. On day 21 (D21), dormant buds began breaking out in the lower part of the stem (trunk) in RG, and 2-3 days later they also broke in the three *MdLAZY1A-W* lines. By D50, each of the 12 trees grew 3-5 straight-upward branches (**Fig. 6B),** prompting termination of the treatment so that the branches could be used for evaluation of branch gravitropic response. On the following day (D0’), the trees were reversed back to their original standing position, effectively turning the branches from an upward position to a nearly horizontal position (**Fig. 6C)**. Branch orientations were measured subsequently on D0’, D3’, D7’, D14’, D21’, D28’ and D42’.

### Subcellular localization of MdLAZY1A

The *MdLAZY1A-S* and *MdLAZY1A-W* coding sequence were cloned into the backbone of vectors pEarleyGate 103, GWB451 and pGreenII 0179 to generate C-terminal GFP and N-terminal YFP fusions, respectively (Hellens et al., 2000; Earley et al., 2006; Nakagawa et al., 2007). The resultant constructs (MdLAZY1A-S::GFP, MdLAZY1A-W::GFP, YFP::MdLAZY1A-S and YFP::MdLAZY1A-W), the plasmids of the plasma membrane aquaporin PIP2A::mCherry and tonoplast aquaporin γ-TIP::mCherry markers (Nelson et al., 2007; Ivanov and Harrison, 2014), and the p19 vector (a common suppressor of post-transcriptional gene silencing) were transiently co-expressed in *N. benthamiana* leaves via *A. tumefaciens* (strains LBA4404 and GV3101 containing the partner plasmid pSoup for the pGreenII backbone)-mediated transformation (Norkunas et al., 2018). Following a 2-3 hour incubation in activation buffer (200 mM MES at pH 5.6, 150 mM MgCl2, and 150 mM acetosyringone), the *Agrobacterium* solution was infiltrated into the abaxial side of *N. benthamiana* leaves of 4-week-old plants. The infiltrated plants were grown for 3-7 days at 23 °C, 50% relative humidity, and 12h light/dark cycles. Transient expression in protoplasts was conducted in *Arabidopsis* Columbia leaf mesophyll protoplasts as detailed previously (Yoo et al., 2007). Following a 30-minute incubation to allow for PEG-mediated protoplast transformation with the desired plasmid(s), protoplasts were washed and re-suspended in 1 ml of W5 solution and incubated in the dark overnight to allow for protein expression. Confocal microscopy was conducted on Zeiss LSM 710, Olympus FV3000, and LeicaTCS SP5 confocal laser scanning microscope systems. Approximately 1-cm^2^ of *N. benthamiana* leaf tissue was excised, and the abaxial side was imaged in water. *Arabidopsis* protoplasts were imaged in W5 solution. Photomicrographs were taken using various combinations of excitation and emission wavelength, including 488 nm and 493-556 nm for EGFP, 514 and 525-550 nm for YFP, 561 nm and 582-650 nm for mCherry, and 633 nm and 652-721 nm for Chlorophyll A. Each image shown is a representative of at least three separate *N. benthamiana* and *Arabidopsis* protoplast transformation experiments. Confocal images were processed using software Fiji (Schindelin et al., 2012).

### Prediction of MdLAZY1A protein features

The deduced protein sequences of MdLAZY1A-S and MdLAZY1A-W were screened with a transmembrane domain (TMD) prediction program TMpred (Hofmann and Stoffel, 1993) and a nuclear localization signal (NLS) identification program cNSL Mapper (Kosugi et al., 2009) to identify putative TMDs and NLSs, respectively. Analysis of the effect of an amino acid substitution on the protein function was conducted using an online tool-Protein Variation Effect Analyzer (PROVEAN) (Choi et al., 2012), and a recommended stringent PROVEAN score with cutoff 4.1 was used to determine if a substitution is deleterious or tolerated.

### Phylogenetic analysis of MdLAZY1A

Protein sequences of LAZY1 orthologs in plants were identified by a BLAST search of the NCBI reference proteins (refseq-protein) databases using AtLAZY1. The cutoff e-value was 0.05. To limit the numbers of accessions, only one species per genera was generally allowed. In addition, isoforms were removed. Alignment of the protein sequences were conducted in groups with T-COFFEE (Notredame et al., 2000). The resultant alignments were combined to generate the final phylogenetic tree using the CLC Genomic workbench software.

### Statistical analysis

One-way analysis of variance (ANOVA) and Tukey-Kramer honestly significant difference (HSD) test were conducted using software package JMP Pro 14 (SAS, Cary, NC). Significance was determined by p<0.05.

### Accession numbers

RNA-seq data of the 12 standard and weeping individuals from the Cheal’s Weeping and Red Jade families can be accessed through NCBI-SRA accession PRJNA738487.

## Supplemental data

Supplemental Figure S1. Schematic representation of the informative recombinants and weeping cultivars as determined by marker genotypes in the *W* region.

Supplemental Figure S2. Screen snapshot of the mapped RNA-seq reads containing mutation c.584T>C (L195P) in the *W* region.

Supplemental Figure S3. Alignment of the genomic sequences of *MdLAZY1A* alleles from weeping (Cheal’s Weeping, Red Jade and Louisa) and standard (Golden Delicious and NY-051) apples.

Supplemental Figure S4. Alignment of the deduced amino acid sequences of MdLAZY1A alleles from weeping (Cheal’s Weeping, Red Jade and Louisa) and standard (Golden Delicious and NY-051) apples.

Supplemental Figure S5. Over-and under-expression of standard allele *MdLAZY1A-S* in apple.

Supplemental Figure S6. Evaluation of gravitropic response of the *MdLAZY1A-W* over-expression lines using growing primary branches.

Supplemental Figure S7. Phylogenetic analysis of 218 LAZY1-like proteins representing 81 plant species in 75 genera.

Supplemental Figure S8. Amino acid sequence alignment of 218 representative LAZY1-like proteins in plants.

Supplemental Figure S9: MdLAZY1A subcellular localization with C-terminal tagged GFP in *Nicotiana benthamiana*.

Supplemental Figure S10. Phenotyping of growth habit in mapping populations segregating for weeping progenies.

Supplemental Figure S11 Diagram for measurement of branch angle and branch tip orientation.

Supplemental Table S1. Segregation of weeping growth in mapping populations derived from Louisa, Cheal’s Weeping and Red Jade.

Supplemental Table S2. List of primers used in study.

Supplemental Table S3. List of genes annotated in the *W* region of 311-kb and their expression levels as determined by RNA-seq analysis on 12 weeping and standard individuals from the families of Cheal’s Weeping and Red Jade.

Supplemental Table S4. RNA-seq reads mapping statistics for the 12 weeping and standard individuals from the families of Cheal’s Weeping and Red Jade.

Supplemental Table S5. Weeping specific SNVs identified in the coding sequence of expressed genes in the W region of 311-kb in the Cheal’s Weeping and Red Jade families.

Supplemental Table S6. Confirmation of allele *MdLAZY1A-S* (carrying CT_584_T) in 28 diverse *Malus* accession of standard growth as determined by their genome sequences (Duan et al 2017).

Supplemental Table S7. Accession number, protein sequence and species information for the 218 LAZY1 orthologs analyzed.

Supplemental Table S8. Frequency of amino acid at a residue position corresponding to L195 in MdLAZY1A in 218 representative LAZY1-like proteins in plants.

Supplemental Table S9. LAZY1 transmembrane domains predicted by TMpred.

## Acknowledgments

We sincerely thank Wlodzimierz Borejsza-Wysocki, Seunghyun Ban, John Phipps, John Calaro, and Selena Lopez for their technical assistance for this study.

